# Altered biological aging-related brain profile in adolescents with autism: A neuroimaging study based on DunedinPACNI

**DOI:** 10.64898/2026.06.19.733477

**Authors:** Javier Rasero, Stefen Beeler-Duden, Peter J. Gianaros, Lauren Kenworthy, Allison Jack, John Darrell Van Horn, Kevin A. Pelphrey

## Abstract

Individuals with Autism Spectrum Disorder (ASD) show increased rates of physical health conditions across major organ systems, some of which are commonly linked with aging, suggesting that body-wide biological profiles may be altered compared to neurotypical controls. Since brain structure associates with peripheral physiology and biomarkers of aging, it may inform understanding of broader biological differences and physical health risks in ASD. Here, we assessed this possibility using DunedinPACNI, a neuroimaging-based model that estimates the pace of longitudinal aging in peripheral organ systems from brain structural information, in 329 adolescents (8-18 years) from the Autism Centers of Excellence network, with replication in an independent age-matched sample of comparable size from the Autism Brain Imaging Data Exchange. Across both datasets and sensitivity analyses, autistic individuals showed significantly larger DunedinPACNI values than controls, consistent with brain structural features that align with patterns associated with a faster pace of biological aging in adults. Group differences were mainly driven by the volume of the 3^rd^ ventricle, cortical thickness of the left entorhinal cortex, grey matter volume of the right entorhinal cortex and grey-to-white matter ratio in the left temporal pole. No association between DunedinPACNI and core autistic traits was found. Our results provide novel evidence for a possible altered brain-body profile in ASD, motivating future studies combining neuroimaging with peripheral biomarkers to better understand the neurobiology of physical health conditions in autism.

## INTRODUCTION

Autism Spectrum Disorder (ASD) is a neurodevelopmental condition characterized by altered communication, social, behavioral and sensory skills, with varying phenotypic expression across individuals, all of which can contribute to challenges related to diagnosis and treatment [1,2]. Some of this heterogeneity can be partially explained by the high co-occurrence of other psychiatric conditions with ASD, including attention-deficit/hyperactivity disorder (ADHD), anxiety disorders, and mood disorders [3,4]. However, less attention is given to the physical health challenges experienced by autistic people, including greater prevalence of or risk for physical health issues such as diabetes, dyslipidemia, and heart and gastrointestinal diseases [5,6]. This multimorbidity suggests that body-level changes (e.g., in tissue and organ function) may be altered in autistic people compared to neurotypical controls and be indicative of processes associated with poorer physical health outcomes, including those typically associated with biological aging.

The autistic brain has been observed to undergo atypical structural changes during childhood and adolescence (6-18 years), particularly an early overgrowth in cerebral volume, combined with accelerated cortical thinning and gyrification and comparable surface area changes [7–10]. These neurodevelopmental differences relate to behavioral and cognitive traits in autism [11,12], suggesting that structural information can be used as multivariate patterns to estimate individuals’ status along their developmental trajectory. While growing evidence links brain structure with peripheral physiology and overall systemic function in early life [13–16], it is yet to be established whether they can be leveraged as a marker of body-wide health information in autistic populations. This reflects, in large part, the scarcity of ASD studies simultaneously collecting markers of biological changes across peripheral systems (e.g., cardiovascular, metabolic, immune) alongside neuroimaging data.

The present study aimed to narrow this gap by examining the degree to which brain structure in autistic adolescents may exhibit patterns consistent with an altered pace of biological aging, via calculation of the DunedinPACNI (Pace of Aging Calculated from Neuroimaging) measure from structural MRI data [17]. We hypothesized that DunedinPACNI would (1) significantly differ between autistic and non-autistic groups and (2) associate with core autistic traits.

## METHODS

### Participants

Our discovery dataset contained 329 individuals comprising neurotypical controls, autistic and unaffected siblings from Wave 1 of the Autism Centers of Excellence (ACE) network. Recruitment and enrollment details for this a multi-site program can be found elsewhere [18]. We also used 286 individuals between age 8-18 years in the Autism Brain Imaging Data Exchange I [19] from four sites that were not part of the ACE network to provide an age-matched replication dataset (Welch’s *t*-test *p*=0.977) of comparable sample size with the discovery set (if excluding unaffected siblings). Descriptive information of both datasets can be found in Table 1.

**Table 1:**
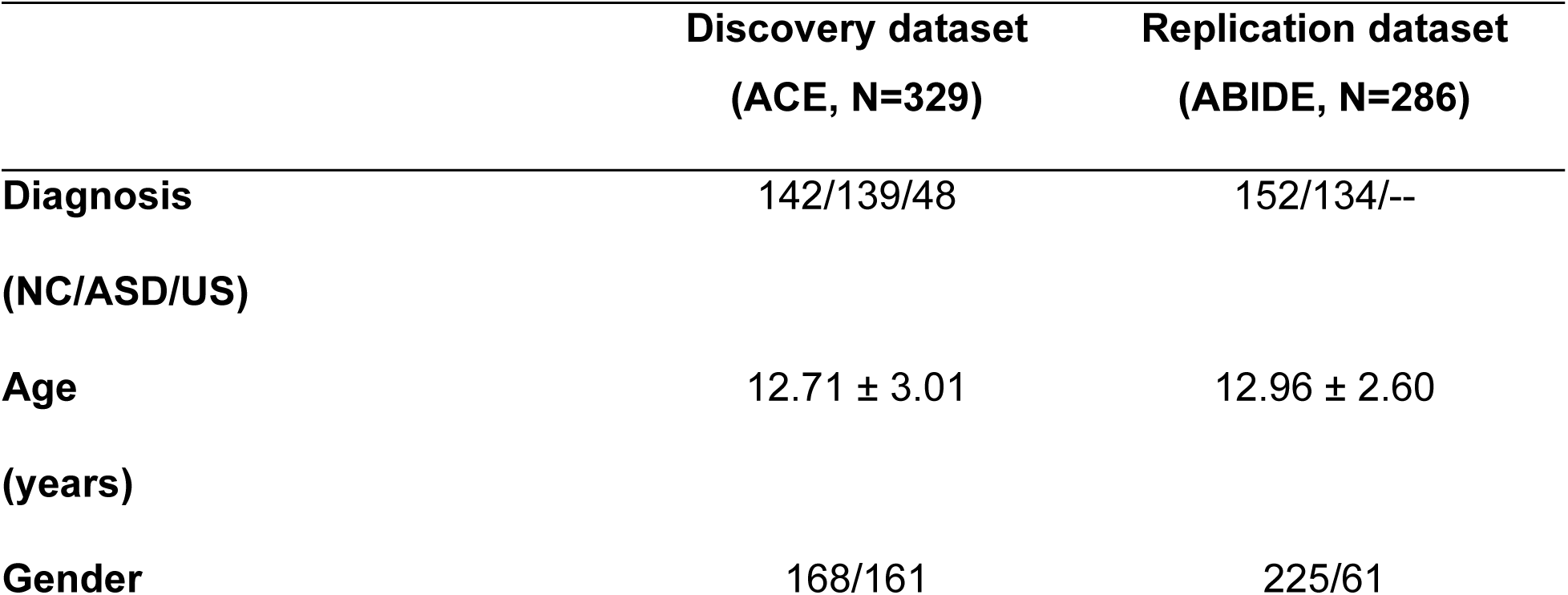
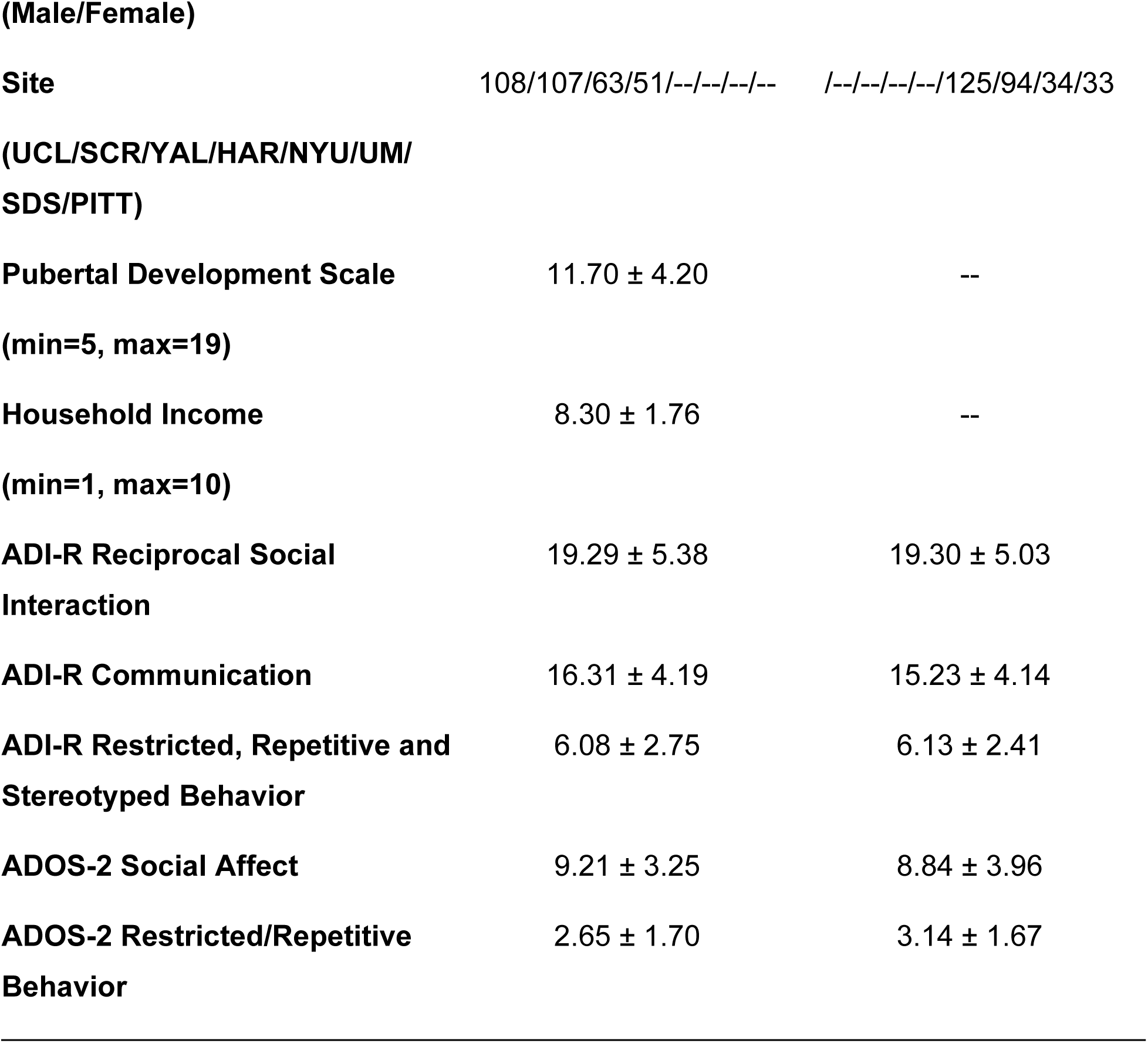
Demographics of our data. NC ≡ Neurotypical Controls; ASD ≡ Autism Spectrum Disorder; US ≡ Unaffected Siblings; UCL≡ University of California (UCLA); SCR ≡ Seattle Children’s/University of Washington; YAL≡ Yale ; HAR ≡ Harvard; NYU ≡ New York University; UM≡University of Michigan; SDS ≡ San Diego State; PITT ≡ Pittsburgh

### MRI preprocessing

T1w images (see [20,19] for image acquisition details) were preprocessed with Freesurfer v6 [21]. CAT12 was also used to extract their image quality rating (IQR) metric, which combines contrast noise, motion-related bias and resolution information, with higher values indicating better quality [22]. We considered only participants with at least sufficient quality (IQR>60). Sensitivity analysis tested higher thresholds.

### DunedinPACNI calculation

A proxy for the pace of biological change was represented through DunedinPACNI [17], a brain-based measure that estimates the aggregated slope of longitudinal change across 19 body-based aging biomarkers from 315 Freesufer ouputs, including cortical thickness, surface area and gray matter volume of the Desikan atlas’ regions [23], gray-white matter signal intensity ratio, subcortical, ventricular, and the bilateral white matter hypo-intensities volumes. To calculate DunedinPACNI, we used the already trained and publicly available algorithm (https://github.com/etw11/DunedinPACNI) and applied it to the same Freesufer outputs of our individuals.

### Statistical analysis

A linear mixed model tested the association of DunedinPACNI with our diagnosis group information, including sex and linear and quadratic age effects as fixed-effect covariates, and site as a random effect. In the discovery dataset, statistical differences across the three groups (neurotypical controls, ASD and unaffected siblings) were evaluated using a Wald test, followed by False Discovery Rate (FDR)-corrected pairwise group comparisons. Brain properties more strongly contributing to observed group differences in estimated DunedinPACNI were assessed by comparing their Shapley Additive Explanations (SHAP) values, a game-theoretic approach quantifying each feature’s contribution to predictions given a trained machine learning model, with positive and negative SHAP values indicating increased and decreased predictions respectively [24]. We also assessed the association of DunedinPACNI with symptom severity across Social, Communication and Behavior subscales of ADI-R [25], and with Social Affect (SA) and Restricted/Repetitive Behavior (RBR) subscales of ADOS-2 [26]. Across all analyses, effect size (β coefficient), standard error (SE), z-statistic and corrected p-values (q-values) were reported.

### Sensitivity analysis

Robustness of the results was tested by (1) restricting analyses to T1w images with satisfactory (IQR>70) and good quality (IQR>80), (2) modeling site information as a fixed effect, and (3) adding separate covariates including Pubertal Development Scale to account for potential pubertal maturation differences [27], and household income as a proxy for socioeconomic status. Finally, to control for the potential influence of clinically significant mental health comorbidities on the observed results, we excluded individuals with T scores greater than 70 in Affective, Anxiety, ADHD, Oppositional Defiant and Conduct Problems from the Child Behavior Checklist. This was done within the discovery set, as the aforementioned covariates were not available in the replication set.

## RESULTS

### Brain structure in autistic individuals exhibits patterns compatible with a faster pace of biological aging

In the discovery set, we found that DunedinPACNI associated with diagnosis (Wald test *χ^2^*(2) =11.137, *p*=0.004). Pairwise comparisons between group categories showed that autistic individuals had significantly larger DunedinPACNI relative to neurotypical controls (β=0.275, SE=0.087, *z*=3.168, *q*=0.005; Figure 1A). This finding was also replicated in an age-matched sample from ABIDE I (β=0.449, SE=0.10, *z*=4.491, *p*<0.001; Figure 1C), despite opposite signs in DunedinPACNI distributions (see Figure 1B), likely attributable to site differences. These group differences were observed in male and female participants separately (Male: β = 0.319, SE=0.124, *z*=2.570, *p*=0.01; Female: β = 0.267, SE= 0.129, *z* =2.062, *p*=0.039), with no evidence of sex moderation (β = −0.039, SE=0.179, *z*=-0.215, *p*=0.83). Autistic individuals also had higher DunedinPACNI values relative to unaffected siblings, although this effect was marginally significant (β = 0.258, SE = 0.128, z = 2.083, *q*=0.056; Figure 1A). No statistical differences between neurotypical controls and unaffected siblings were found (β = −0.017, SE = 0.124, *z* = −0.137, *q*=0.891; Figure 1A).

**Figure 1.**
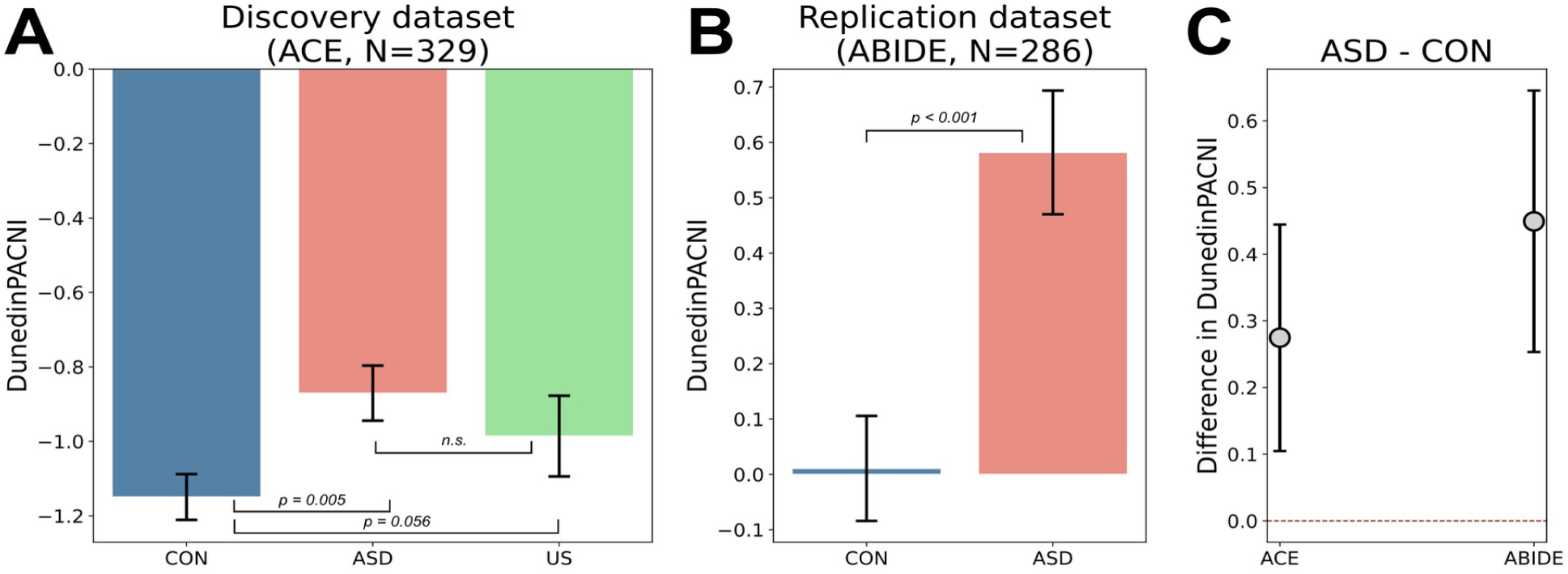
Group differences in DunedinPACNI, representing the pace of biological aging-like change as predicted from MRI scans, in A) the discovery set (ACE) and B) the replication set (ABIDE). C) Across both datasets, the estimated difference and 95% confidence intervals in DunedinPACNI between autism and neurotypical control groups. CON≡Neurotypical Controls; ASD≡Autism; US≡Unaffected siblings

### Core autism traits are not associated with DunedinPACNI

There was no significant association between DunedinPACNI and ADI-R subscale scores in the discovery sample (Social: β=0.005, SE=0.013, z=0.396, p=0.692; Communication: β= 0.012, SE=0.017, z=0.688, p=0.491, Behavior: β= 0.005, SE=0.028, z=0.168, p=0.866) and ADOS-2 (SA: β=-0.025, SE=0.022, *z*=-1.100, *p*=0.271; RBR: β=0.010, SE=0.041, *z*=0.242, *p*=0.809). Similar null associations were observed in the replication dataset.

### Brain properties most contributing to group differences

Differences in SHAP values between autistic and neurotypical individuals in the discovery dataset for all features with nonzero regression coefficients in the DunedinPACNI algorithm are shown in Figure 2A. Only four features survived FDR correction, including significantly larger values for autistic individuals relative to neurotypicals in volume of the 3^rd^ Ventricle (*t(279)*=3.633, *q*=0.033), cortical thickness of the left entorhinal cortex (*t(279)*=3.407, *q*=0.033) and grey matter volume of the right entorhinal cortex, (*t(279)*=3.26, *q*=0.033), as well as significantly lower values in grey-to-white matter ratio in the left temporal pole (*t(279)*=-3.245, *q*=0.033)). These specific properties were not the largest contributors within the algorithm for DunedinPACNI estimation, meaning that the observed group differences did not simply reflect dominant features in this model. There were also no significant linear or quadratic age-by-group interaction effects for these features.

**Figure 2.**
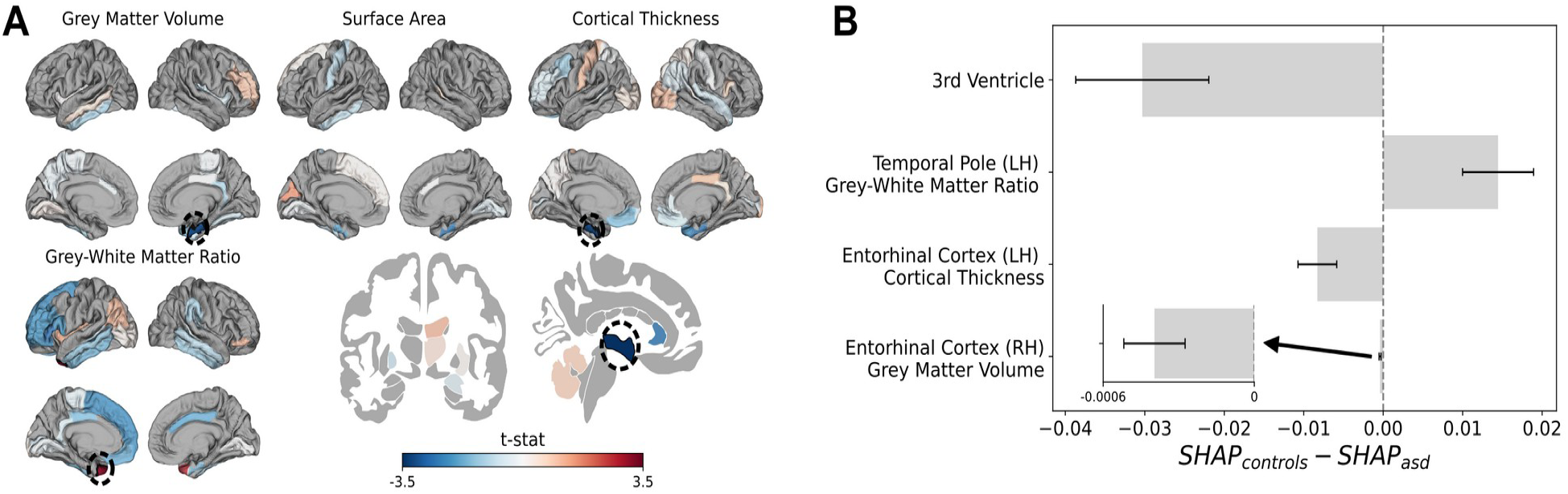
A) Within the discovery dataset, statistical maps of autistic versus neurotypical differences in SHAP values for the brain structural features used by the deployed DunedinPACNI algorithm. Dashed circles highlight those features that were significant after FDR correction. B) Distribution of group differences in SHAP values for these features.

### Sensitivity Analysis

Our results remained significant after imposing higher imaging quality thresholds (IQR>70: β = 0.195, SE= 0.078, z = 2.507, p=0.012; IQR>80: β = 0.287, SE= 0.103, *z* = 2.787, *p*=0.005) and when modeling site information as a fixed effect (β = 0.272, SE= 0.089, *z* = 3.064, *p*=0.002). Results were also robust when accounting for differences in pubertal maturation (β = 0.267, SE= 0.097, *z* = 2.749, *p*=0.006) and household income (β = 0.407, SE= 0.109, z = 3.748, p<0.001). Finally, group differences held when focusing on individuals without severe concurrent mental health comorbidities (β = 0.246, SE= 0.102, *z* = 2.407, *p*=0.016).

## DISCUSSION

Here, we tested whether brain structure in ASD during childhood and adolescence yields estimates compatible with an altered pace of body-wide biological aging, as indexed by DunedinPACNI, a neuroimaging-based measure previously developed in adults. Our results using the ACE dataset and further replicated in an age-matched cohort from ABIDE supported this hypothesis; specifically, autistic individuals had significantly higher DunedinPACNI values, suggesting that their brain patterns may be consistent with a faster pace of biological aging compared to neurotypical controls. Notably, their DunedinPACNI values did not appear to associate with core autism features. These results were robust to data quality thresholds, site effects modeling, adjustment for pubertal maturation and socioeconomic factors, and exclusion of individuals with clinically significant mental health symptoms.

Although still scarce, prior work has reported alterations in autistic youth across several biological systems included in the original DunedinPACNI study. Specifically, autistic adolescents are more likely to develop periodontitis and other dental-related diseases [28,29], rank in higher BMI percentiles (especially in early adolescence) [30], exhibit distinct lipid and apolipoprotein profiles [31], have higher systemic inflammation levels [32,33], and show evidence of altered respiratory-autonomic functioning [34,35]. Therefore, our results appear to align with this growing evidence, but here through brain structure profiles resembling those previously shown to capture peripheral, body-level changes in adults.

Our results indicate that brain-related biological change may not vary systematically across the spectrum, since we found no significant correlations between DunedinPACNI and standard autism diagnostic scales (ADOS and ADI-R). To date, only one study has investigated this question, providing evidence for an association between autistic traits and a faster pace of physiological change [36]. However, this was based on an adult population, which may be more sensitive to aging effects, and employed different assessment tools compared to our study. Thus, we cannot rule out the possibility that similar associations may emerge when using other measures of autistic traits.

The strongest individual brain properties underlying the observed group differences also seem to converge with prior reports of atypical ventricular morphology, cortical thickness, grey-matter morphometry, and grey–white matter boundary contrast in autism. One may then wonder whether this means that an approach such as brain age gap [37,38], which is known to be altered in brain disorders during development [39], would have captured the same effect. However, we found no group differences in the relationship between our reported brain properties and age (i.e., age-by-group interactions), which may indicate that their contribution to brain age prediction would be small. Furthermore, moderate correlations between brain models of chronological age and biological aging have been reported [17,40], potentially supporting their complementary value for characterizing brain disorders.

The main limitation of the present study is the application of DunedinPACNI to our cohort, as this measure was calculated from a model using biomarkers of biological aging and cross-sectional MRI data in midlife adulthood. Therefore, our results were interpreted as capturing biological aging-like effects rather than actual biological developmental changes. Moreover, we focused throughout on relative differences between groups, who were matched by age, rather than on absolute values, which should be interpreted with caution. Future studies should aim to create similar models in younger populations by collecting the same or comparable biomarkers as those in the DunedinPACNI study to estimate the pace of change across multiorgan systems and linking them with neuroimaging data.

Despite this limitation, we showed replicable significant group differences that were also robust to multiple analytical choices. We therefore hope the present study encourages the collection of body-wide biomarkers alongside brain data in future ASD research efforts, as this may shed light on the neurobiology underlying poor physical health outcomes in this neurodivergent population.

## ACKNOWLEDGEMENTS

The authors sincerely thank Ethan Whitman for his feedback and guidance in interpreting the DunedinPACNI-related results.

## AVAILABILITY OF DATA AND MATERIALS

The raw data for the discovery dataset (ACE network) can be downloaded from the National Institute of Mental Health Data Archive (collection #2021). The raw data for the replication dataset are part of ABIDE, which is openly available.

## CONFLICT OF INTERESTS

The authors declare that they have no competing interests.

## Notes

### Competing Interest Statement

The authors have declared no competing interest.

